## Additional File 1 for "Comparative Transcriptomic Signature of the Simulated Microgravity Response in *Caenorhabditis elegans*"

### Supplementary Figures

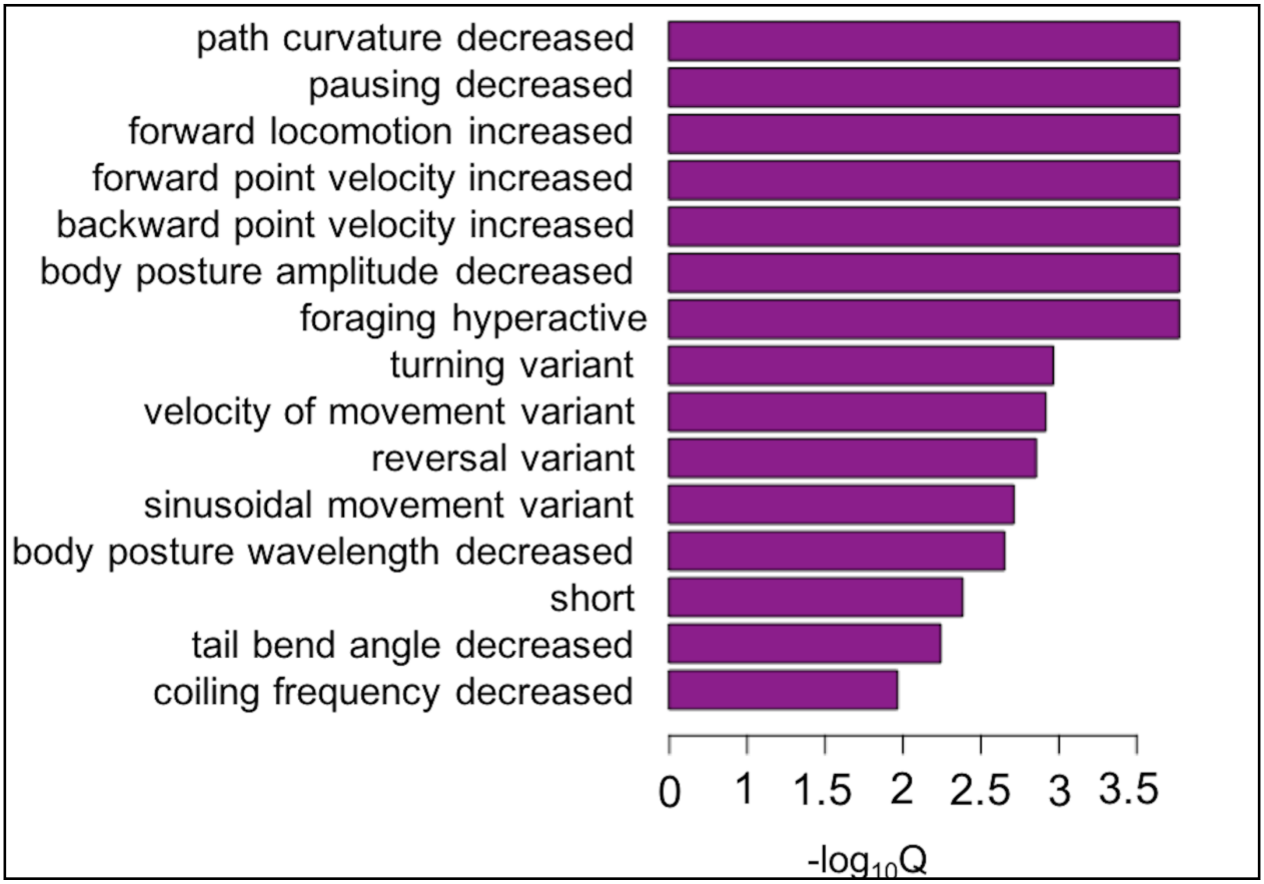


**Fig. S1:** Phenotype enrichment analysis of the DEG neuropeptide signaling genes indicate a movement variation under simulated microgravity.

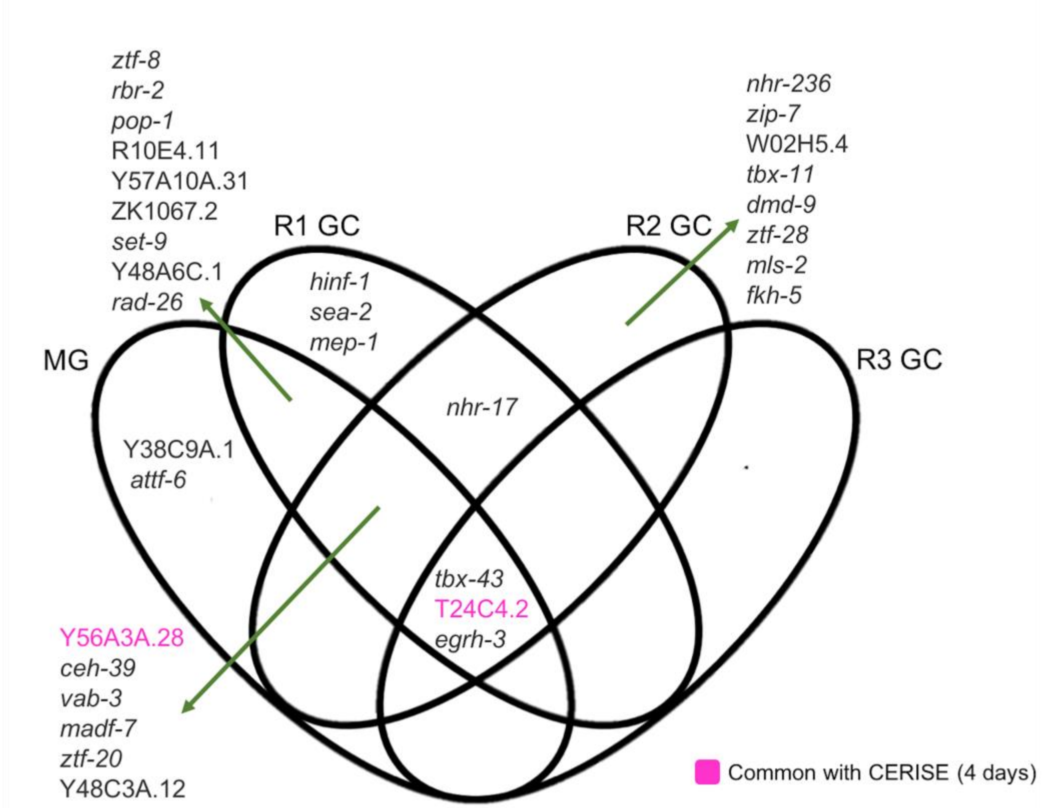


**Fig. S2:** Putative transcription factor genes that are upregulated under simulated microgravity (MG), four days (R1 GC), eight days (R2 GC), and twelve days (R3 GC) after return to ground conditions. Y56A3A.28 and T24C4.2 are also induced in four-day CERISE.

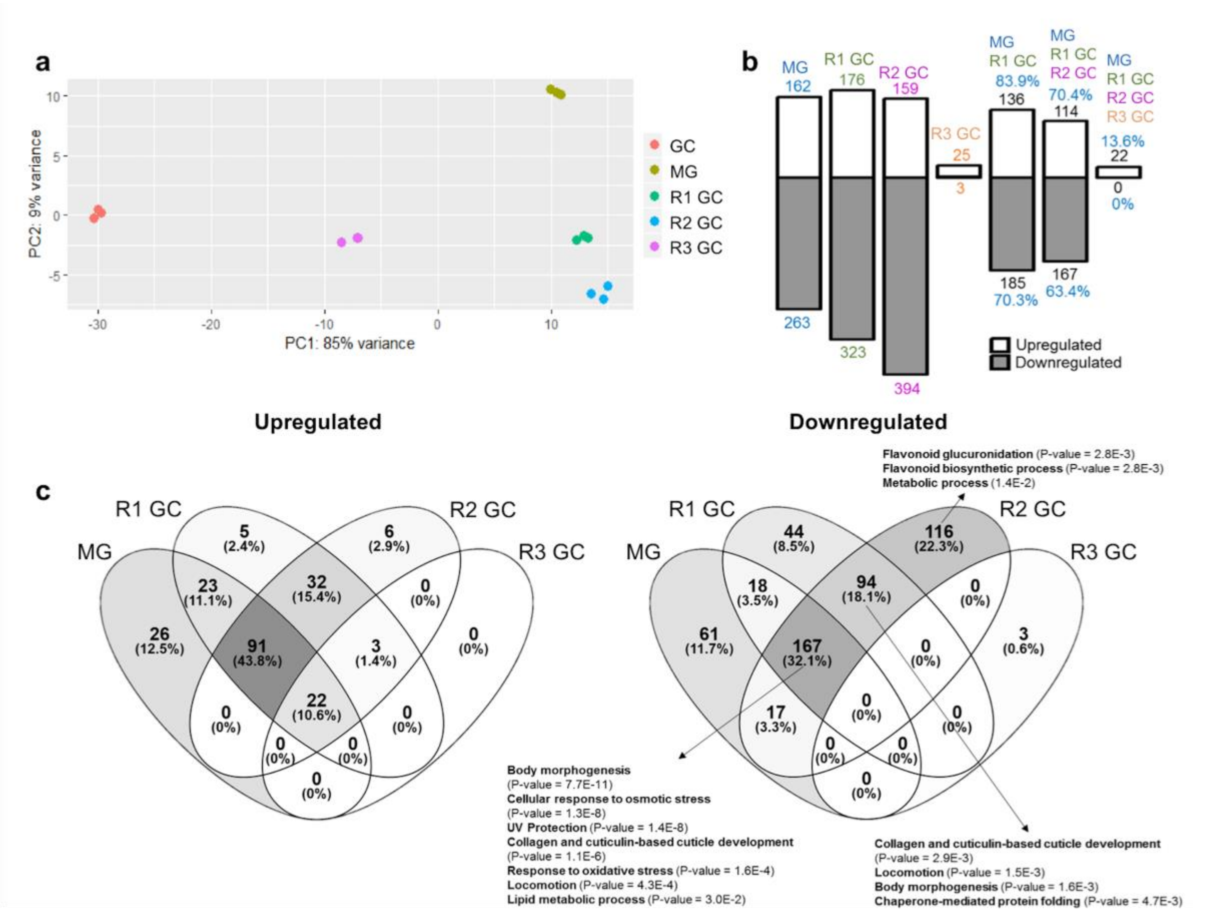


**Fig. S3:** RNA-seq data analysis with DESeq2 **(A)** PCA analysis of the RNA-seq data from the ground control (GC), simulated microgravity (MG), and four-, eight-, and twelve-day after return to ground conditions. **(B)** The number differentially expressed genes under simulated microgravity and the transmission of the differential expression after return to ground conditions from the analysis with DESeq2. **(C)** Categorization of the upregulated (left) and downregulated (right) genes in comparison to the ground control animals, and the enriched gene ontology terms assigned to them from the analysis wirh DESeq2.

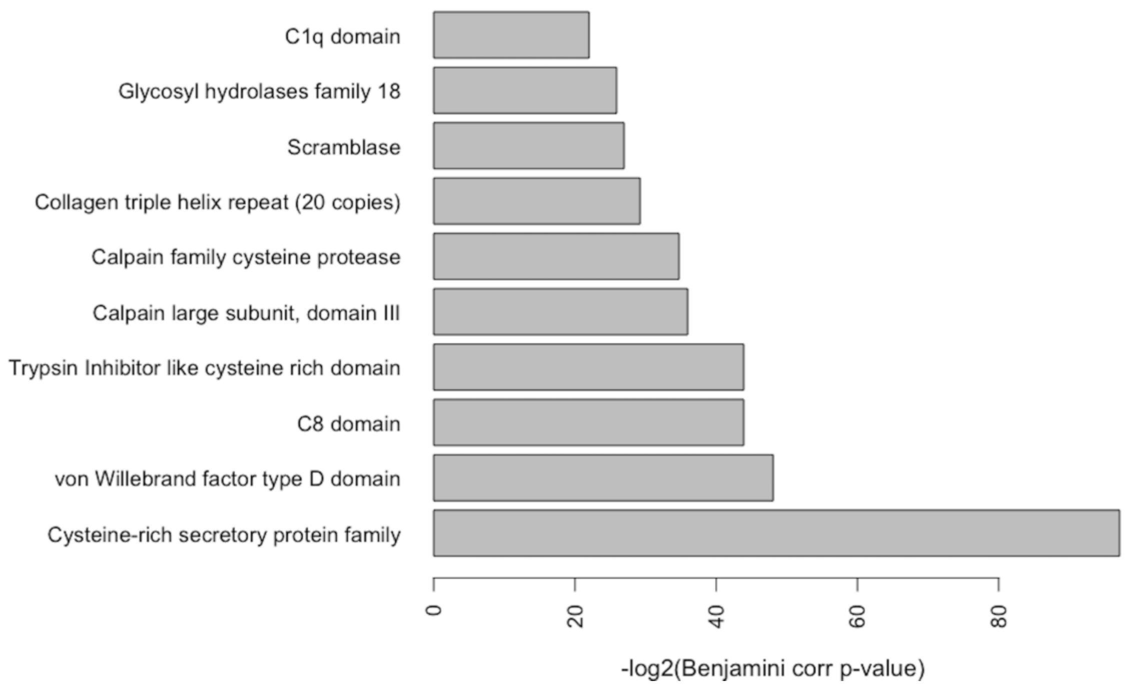


**Fig. S4:** Domain enrichment of the human orthologs of the gravitome.

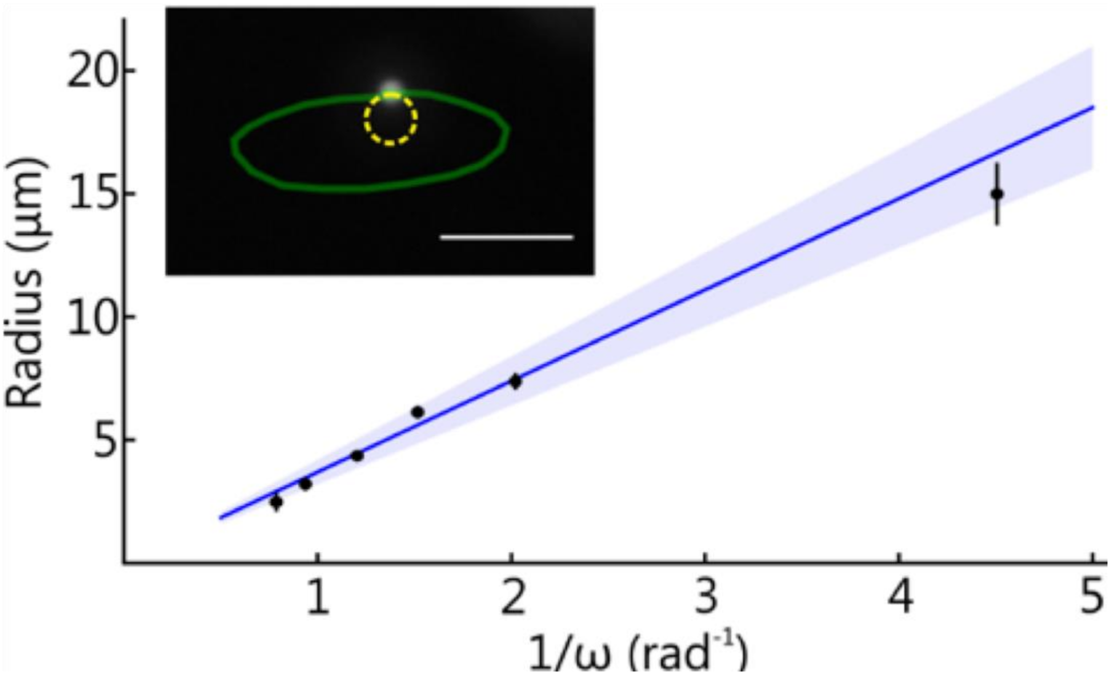


**Fig. S5:** The mean radius of the motion of a suspended melamine-formaldehyde microsphere versus angular velocity of the clinostat (*black*) compared to the expected radius (*blue*). The error bars represent the standard error of measurement. The shaded area represents the expected radii for terminal velocities within one standard deviation of the mean. The inset shows the orbit of a single microsphere (ω = 1 rad/sec) overlaid on the uncorrected trajectory (*green, solid*) and trajectory corrected for mechanical noise (*yellow, dotted*) using fiducials. The scale bar in the inset is 20 μm.

### Supplementary Dataset Legends

**Supplementary Dataset S1:** Sequencing read information for the RNA-seq libraries.

**Supplementary Dataset S2:** Venn categorized DEGs represented in Fig. 2d, e. The data were analyzed with the Tuxedo pipeline.

**Supplementary Dataset S3:** Upregulation or downregulation of DAF-16 induced and suppressed genes.

**Supplementary Dataset S4:** Different acyl-chain and total ceramide levels.

**Supplementary Dataset S5:** Venn categorized DEGs from the DESeq2 analysis of the RNA-seq data.

**Supplementary Dataset S6:** LogFC comparisons of the gravitome from our data, four-day CERISE and PRJNA146465.

**Supplementary Dataset S7:** Human orthologs of the gravitome.
